# Host-specific epibiomes of distinct *Acropora cervicornis* genotypes persist after field transplantation

**DOI:** 10.1101/2021.06.25.449961

**Authors:** Emily G. Aguirre, Wyatt C. Million, Erich Bartels, Cory J. Krediet, Carly D. Kenkel

## Abstract

Microbiome studies across taxa have established the influence of host genotype on microbial recruitment and maintenance. However, research exploring host-specific epibionts in scleractinian corals is scant and the influence of intraspecific differences across environments remains unclear. Here, we studied the epibiome of ten *Acropora cervicornis* genotypes to investigate the relative roles of host genotype and environment in structuring the epibiome. Coral mucus was sampled in a common garden nursery from replicate ramets of distinct genotypes (T_0_). Coral fragment replicates (n=3) of each genotype were then transplanted to nine different field sites in the Lower Florida Keys and mucus was again sampled one year later from surviving ramets (T_12_). 16S rRNA amplicon sequencing was used to assess microbial composition, richness, and beta-diversity. The most abundant and consistent amplicon sequencing variants (ASVs) in all samples belonged to Fokiniaceae (MD3-55 genus) and Cyanobacteria (*Synechococccus*). The abundances of these bacterial taxa varied consistently between genotypes whereas neither the composition nor taxonomic abundance were significantly different among field sites. Interestingly, several high MD3-55 hosting genotypes showed rapid diversification and an increase in MD3-55 following transplantation. Overall, our results indicate healthy *A. cervicornis* genotypes retain distinct epibiome signatures through time, suggesting a strong host component. Lastly, our results show that differences in MD3-55 abundances can be consistently detected in the epibiome of distinct host-genotypes of *A. cervicornis*. As this organism (sensu *Aquarickettsia rohweri*) has been implicated as a marker of disease resistance, this finding reinforces the potential use of microbial indicators in reef restoration efforts via non-invasive surface/mucus sampling.

## Introduction

Microbiome studies across taxa link host specificity to distinct microbial ecotypes, most famously in humans (Kolde et al. 2018; Lynch and Hsiao 2019), plants (Wagner et al. 2016), insects (Vogel and Moran 2011) and recently in acroporid corals (Glasl et al. 2019). Mounting evidence has established the diversity and importance of bacterial associates in corals (Bourne et al. 2016; Sweet and Bulling 2017; van Oppen and Blackall 2019). However, dissecting the influence of host genotype on microbial recruitment and maintenance in corals remains a challenge due to holobiont (host and its collective microbial associates) diversity (Blackall et al. 2015) and microhabitat niche distinctions (e.g. surface mucus, tissue, and skeleton in scleractinian coral) (Apprill et al. 2016). Despite these challenges, investigations into host-specific bacterial associates in corals can help partition the effect of genotype on the microbiome and its relationship to holobiont fitness and disease.

Acroporid coral microbiomes are known to vary among conspecific hosts but knowledge about the combined effect of environment and genotype on the stability of microbiomes (Glasl et al. 2019; Marchioro et al. 2020; Miller et al. 2020) and its link to holobiont resilience is limited. Previously, (Wright et al. 2017) challenged *Acropora millepora* genotypes with pathogenic *Vibrio* spp. to determine if coral disease was a response to an etiological agent or to a weakened holobiont. Disease resistant genotypes were largely unaffected by *Vibrio* spp., and gene expression resembled that of healthy non-inoculated corals, suggesting that coral disease results from an unfavorable combination of genotype and environment. Similarly, *A. cervicornis* genotypes exhibit distinct tissue microbial signatures (Klinges et al. 2020; Miller et al. 2020) differentiated by varying abundances of a Rickettsiales coral symbiont, and presumed to be an indicator of disease susceptibility during thermal stress (Klinges et al. 2020). The influence of the environment on distinct *A. tenuis* genotypes has also been studied in experimental manipulations, suggesting distinct host-genotype specific microbial composition, irrespective of single stress or combined stress treatments (e.g., reduced salinity, thermal stress, elevated pCO_2_ and presence of a macroalgae competitor (Glasl et al. 2019). Taken together, this suggests that acroporid genotypes exhibit unique microbiome signatures, but it is unclear if these geno-typespecific microbiomes are maintained through time in natural reef environments. Investigating the influence of host-specificity and environment is foundational to unraveling microbially-mediated processes underpinning the maintenance of holobiont health in conspecific hosts.

The coral surface mucus is an ideal microhabitat to explore genotype and environment dynamics due to its function as a defensive barrier between coral epithelia and the environment, and its putative role in preventing/causing disease (Sutherland et al. 2004; Brown and Bythell 2005; Krediet et al. 2013). Coral mucus composition varies among coral species (Meikle et al. 1988), but generally it is a polysaccharide protein lipid complex, and can be viewed as a secretory product with multiple functions (Crossland 1987; Coffroth 1990; Wild et al. 2004). Because of its rich organic composition, mucus hosts the highest bacterial diversity (Garren and Azam 2010) and contributes to nutrient cycling in the holobiont (Wild et al. 2004). Surface and mucus microbiomes (epibiomes) are presumed to be influenced by the surrounding environment (Kooperman et al. 2007; Pollock et al. 2014; McDevitt-Irwin et al. 2017). Therefore, thorough epibiome characterizations in coral holobionts are pivotal as reefs respond to oceanic changes due to climate change (Ritchie 2006). Recent research exploring the intersection between environment and acroporid epibiomes yielded novel insights in *A. tenuis* and *A.millepora*, showing that epibiomes were very different from tissue microbiomes, and shared a similar microbial composition as the surrounding seawater (Marchioro et al. 2020). However, host-genotype responses, in the epibiomes of acroporids, to variable environments remain under-explored (Marchioro et al. 2020; Miller et al. 2020).

A fundamental understanding of genotype and environment dynamics in coral epibiomes can aid restoration efforts since bacterial communities are implicated in coral health and holobiont resistance (Krediet et al. 2013; Peixoto et al. 2017; Rosado et al. 2019; van Oppen and Blackall 2019). Here, we aimed at addressing this knowledge gap for the staghorn coral, *Acropora cervicornis*, an ecologically relevant and endangered Caribbean coral. Our goal was to determine the extent of host genotype, environment or the combination in the maintenance of epibiomes over time. To do this, we assessed the environmental response of the epibiome among and within *A. cervicornis* genotypes following transplantation to novel reef environments. We found that *A. cervicornis* genotypes exhibited distinct epibiome signatures, driven by abundance of MD3-55, a ubiquitous Rickettsiales bacterial symbiont. Alpha-diversity and beta-diversity of bacterial communities shifted slightly in surviving outplants, regardless of the ultimate reef site, following one year of field transplantation. We also observed an increase of MD3-55 in several high MD3-55 hosting genotypes. Despite changes in MD3-55 relative abundance, *A. cervicornis* genotypes retained their unique microbial signatures, suggesting a strong host component contributes to epibiome community composition.

## Methods

### Study Overview

We sampled mucus from replicate fragments (n=30) of ten known coral genotypes from Mote Marine Laboratory’s *in situ* nursery (24° 33’ 45.288” N, 81° 24’ 0.288” W) in April 2018. Fragments from all ten genotypes were propagated long-term on mid-water structures (coral “trees”) for at least 5 years, and then mounted on concrete discs and attached to benthic modules in preparation for transplantation at least two weeks prior to mucus sample collection. The samples were prepared for 16S rRNA amplicon sequencing to assess alpha-diversity between fragments of the same genet and beta-diversity between genotypes (T_0_). Following nursery sampling, replicate ramets (n=3) of each genet were transplanted to nine different field sites (Table S1) in the Lower Florida Keys in April 2018. Concrete discs were attached to the reef substrate with marine epoxy. Metal tags were used to identify colonies for future surveys. We returned to the sites in April 2019 to again collect mucus samples to assess changes in the epibiotic microbiome of surviving ramets (T_12_).

### Mucus Sampling and 16S rRNA Sequencing

Epibiome samples were obtained by agitating the coral surface with a 10 mL syringe to stimulate mucus production (Ritchie 2006). Mucus samples were then transferred to 15 mL Eppendorf tubes and frozen at −20°C until processing. Background filtered seawater (FSW) controls were obtained by filtering duplicate 1L seawater samples through a 1.0μm pore-size, 47mm polycarbonate filter (Whatman International, Ltd., England), and size fractionated through a 0.2 μm pore-size filter, to capture bacteria between 1-0.2μm and frozen at −20°C until processing.

The samples were prepared for processing by thawing and centrifuging for 30 minutes, where only the bottom, heavier fraction (~2-3 mL) containing the mucus was concentrated and the seawater supernatant was discarded. DNA was extracted from all samples using the DNEasy PowerBiofilm Kit (Qiagen, Hilden, Germany), followed by targeted amplification of the V4 region of the 16S rRNA gene using the Earth Microbiome Project protocols (Thompson et al. 2017) along with the 515F-806R primer set (Caporaso et al. 2011; Parada et al. 2016). Unique Illumina barcodes were incorporated in a second round of PCR and samples were pooled in equimolar amounts for sequencing of paired 250-bp reads on the Illumina MiSeq v2 PE 250 platform (Admera Health, USA).

### Bioinformatic and Data Analysis

We were able to amplify and successfully sequence 128 samples from the initial timepoint (T_0_) and 112 samples from the one-year timepoint (T_12_). Resulting paired-end reads were demultiplexed and quality checked using FastQC (Andrews 2010). Amplicon sequencing variants (ASVs) were called using DADA2 (Callahan et al. 2016) in R (R Core Team 2020), using the default standard filtering parameters (truncLen=c(240,160),maxN=0, maxEE=c(2,2), truncQ=2, rm.phix=TRUE, compress=TRUE, multithread=TRUE) and constructed into a sequence table. Taxonomy was assigned using the naive Bayesian classifier method (Wang et al. 2007) in conjunction with the Silva SSU training data for DADA2, version 138 (DOI 10.5281/zenodo.3731174). Statistical analyses and visualizations were conducted in R (R Core Team, 2020). The arrayQualityMetrics and DESeq2 R packages (Kauffmann et al. 2009; Love et al. 2014) were used to screen for outlier samples. The compositional nature of data generated by high-throughput sequencing requires normalization techniques to transform the data into a symmetrical dataset (Gloor et al. 2017; Weiss et al. 2017), therefore we rarefied to an even read depth (5,000 reads), and samples <5,000 reads were discarded (Table S2). After rarefaction, our dataset contained 8,026,016 reads.

Sequence removal of mitochondria and eukaryotes, rarefaction, relative abundance visualizations, alpha-diversity (Shannon’s Index) and beta-diversity plots were generated using Phyloseq (McMurdie and Holmes 2013) and ggplot2 (Wickham 2016). To determine consistent bacterial taxa, we conducted core microbiome analysis in the Microbiome package (Lahti and Shetty 2019) using a detection limit of 0.01% in >60% of the samples (prevalence threshold). Cyanobiaceae (Synechococcales) and Fokiniaceae (Rickettsiales order) were identified as the two dominant amplicon sequencing variants in all T_0_ coral samples. We queried our Fokiniaceae ASVs with that of two published sequences of coral-associated Fokiniaceae genera: MD3-55 in the NCBI database and a full-length 16S rRNA sequence of Candidatus *Aquarickettsia rohweri* (Klinges et al. 2019). The former were also queried with 16S rRNA gene sequences of the Rickettsiales order (12 families) and one non-Rickettsiales representative in Alphaproteobacteria was chosen as the outgroup (*Caulobacter mirabilis*). Phylogenetic analysis was carried out by aligning all sequences using the *MUSCLE* algorithm (Edgar 2004). The aligned sequences were used to construct a maximum likelihood phylogeny with ultrafast bootstrap (1000 bootstrap replicates) using *IQTREE* (Kalyaanamoorthy et al. 2017; Minh et al. 2020) and visualized on the Interactive Tree of Life interface (https://itol.embl.de/itol.cgi).

We conducted multivariate analyses to test observed dissimilarities between (beta-diversity) microbial communities of genotypes hosting low abundances of MD3-55 versus genotypes hosting high abundances of MD3-55, across timepoints using the vegan package (Oksanen et al. 2019). Differences in groups were visualized by principal coordinate analysis (PCoA) using the weighted-Unifrac metric (Fig. 1) and statistical differences were determined using the non-parametric tests, analysis of similarities (ANOSIM) and permutational multivariate analysis of variance (PERMANOVA). First, homogeneity of group dispersions was tested in pairwise comparisons, followed by the appropriate non-parametric test. If uneven group dispersion was prevalent, the ANOSIM test was applied. Pairwise comparisons with even group homogeneity were tested using PERMANOVA. ASVs with <5 counts in at least 1 of the samples were excluded from relative abundance visualizations, beta-diversity, and multivariate analyses.

**Figure 1.**
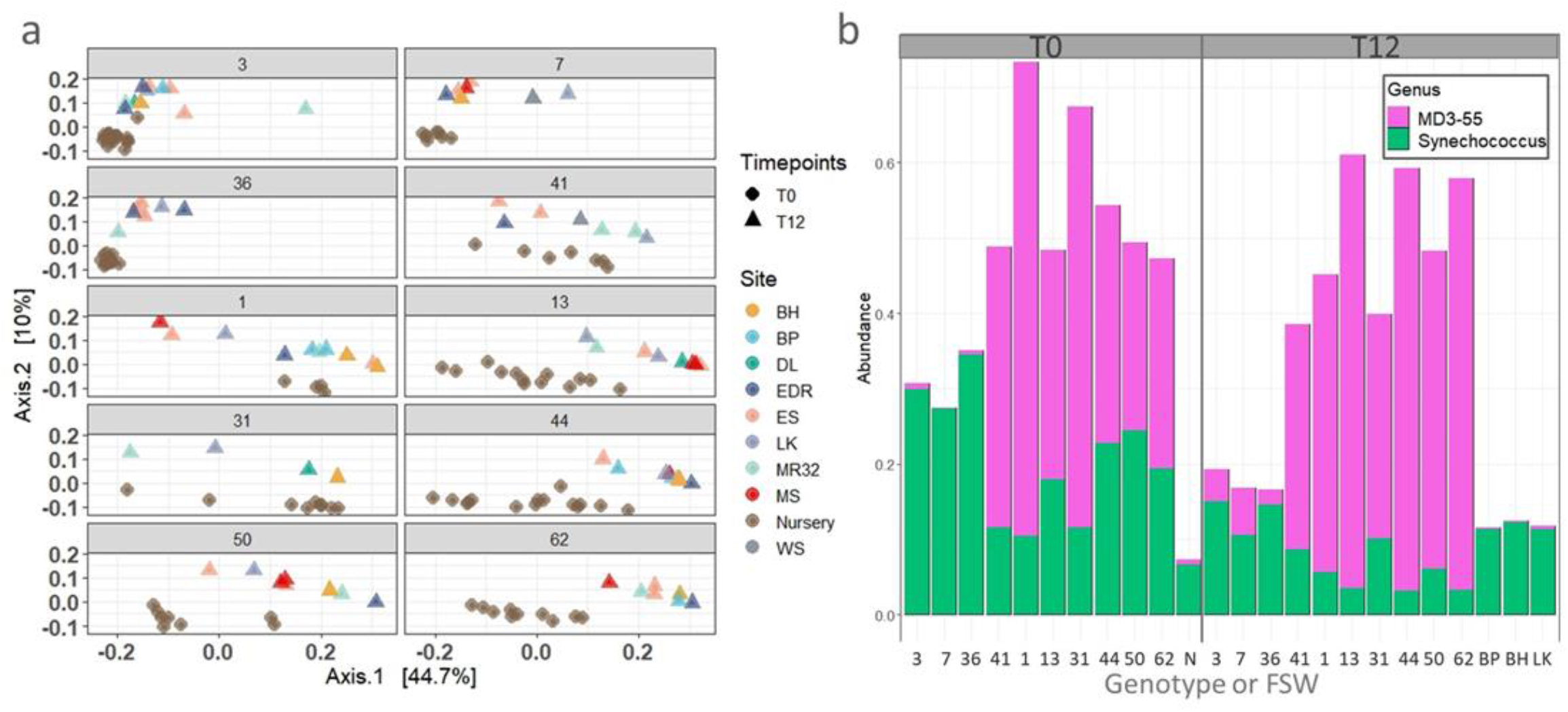
Beta-diversity and most abundant taxa present in all samples. (a) Principal coordinates analysis (PCoA) on a weighted-Unifrac metric, used to visualize differences between *Acropora cervicornis* genotypes (G3, G7, G36, G4l, Gl, G13, G3l, G44. G5O, G62), site (colors) and timepoint (circles= T_0_ and triangles=T_12_). (b) Relative abundance of the top 100 ASVs in the dataset: MD3-55 (Fokiniaceae family, Rickettsiales order) and *Synechococcus* CC99O2 (Synechococcaceae family. Cyanobacteria class) by genotype and background FSW (labeled as N: Nursery, BP: Big Pine, BH: Bahia Honda and LK: Looe Key).

## Results

### *A. cervicornis* epibiome composition

We obtained 9,695,036 reads from 240 samples (coral and seawater controls). A total of 8,104,340 reads and 21,554 ASVs (Table S3) were retained after filtering for mitochondria, chloroplast and protistan variants. Samples containing less than 5,000 reads were removed and the remaining 191 samples (Table S3) were subsequently rarefied to the minimum even read depth of 5,000 reads, resulting in 16,850 ASVs (Table S2).

The predominant phyla identified in the dataset were Proteobacteria (56%), Cyanobacteria (26%), Bacteroidetes (10%), Actinobacteria (4%) and Spirochaetes (1%). All other phyla were present in abundances <1% (Fig. S1). Alpha-diversity (calculated using all ASVs) was high in all genotypes, but highest in FSW samples (Fig. S2). Alpha-diversity also increased when corals were transplanted to field sites (T_12_) resulting in statistical differences between the two timepoints (Wilcoxon, p = 0.03, Table S4). Differences in beta-diversity were observed between the nursery FSW samples and nursery coral samples (ANOSIM R= 0.786, p = 0.001). However, we were only able to compare the bacterial composition of coral mucus and background water samples for two sites in the field (T_12_). Beta dispersion tests for FSW samples from Bahia Honda (BH, n=2) and Big Pine (BP, n=2) sites were conducted against the coral samples from their respective sites and found to be evenly dispersed and significantly different for both BH (BETADISPER, p=0.9, PERMANOVA, p= 0.02) and BP (BETADISPER, p=0.7; PERMANOVA, p=0.03) (Table S5). Additionally, no significant differences in beta-diversity were detected in pairwise comparisons of background seawater controls (nursery, BP and BH, Table S5).

Alpha-diversity assessments of the epibiome, using Shannon’s index, showed differences among genotypes (ANOVA, F = 3.5, *p* = 0.0006). Post-hoc analyses (Tukey multiple comparison of means) detected three significant differences among pairwise comparisons: genotypes G44-G36 (adjusted p < 0.02), G44-G41 (adjusted p < 0.04) and G7-G44 (adjusted p < 0.04) (Table S6).

### *A. cervicornis* epibiomes exhibit distinct genotype signatures

*A. cervicornis* genotypes reared in a common garden nursery environment (T_0_) exhibited distinct signatures in the overall epibiome that largely persisted, even after one year of transplantation to distinct field environments (T_12_) (Fig. 1a). The top 100 most abundant ASVs that were present in the coral samples were *Synechococcus* (Cyanobacteria) and MD3-55 (Rickettsiales) (Fig.1b), with genotypes 3, 7 and 36 hosting low abundances of MD3-55 and the remaining genotypes hosting high MD3-55 abundances. Low MD3-55 hosts (G3, G7 and G36) differed from high MD3-55 hosts (all other genotypes, ANOSIM *R* = 0.627, *p* < 0.001) but microbial community composition was similar and not statistically different when MD3-55 ASVs were removed (ANOSIM, *R* = 0.016, *p* = 0.3).

### *A. cervicornis* epibiomes exhibit temporal changes but not among-site differences

Temporal changes in the epibiome composition of the ten genotypes were also assessed, and while the groups (T_0_ versus T_12_) were evenly dispersed (BETADISPER, p=0.294) beta-diversity between the two sampling timepoints was significantly different (PERMANOVA, p=0.001, Fig.1a). *A. cervicornis* epibiomes did not differ among transplant sites in terms of alpha-diversity (Kruskal Wallis test, p=0.95, Table S4) or composition at the phylum level (Fig. S3).

### Rickettsiales and Cyanobacteria are consistent members of the *A. cervicornis* epibiome

To identify taxa that were consistently present in the epibiome across sampling timepoints (T_0_ and T_12_), we conducted core microbiota analysis using the microbiome package in R on the filtered dataset. The prevalence threshold was set to 60% of the samples and a moderate detection limit of 0.002% was applied. Out of the 6,966 analyzed ASVs, only 190 ASVs were consistent in >60% of the samples (Fig. 2). The detected 190 ASVs consisted of 2 bacterial taxa, MD3-55 (Rickettsiales) and Synechococcus CC9902 (Cyanobacteria) (Fig.2).

**Figure 2.**
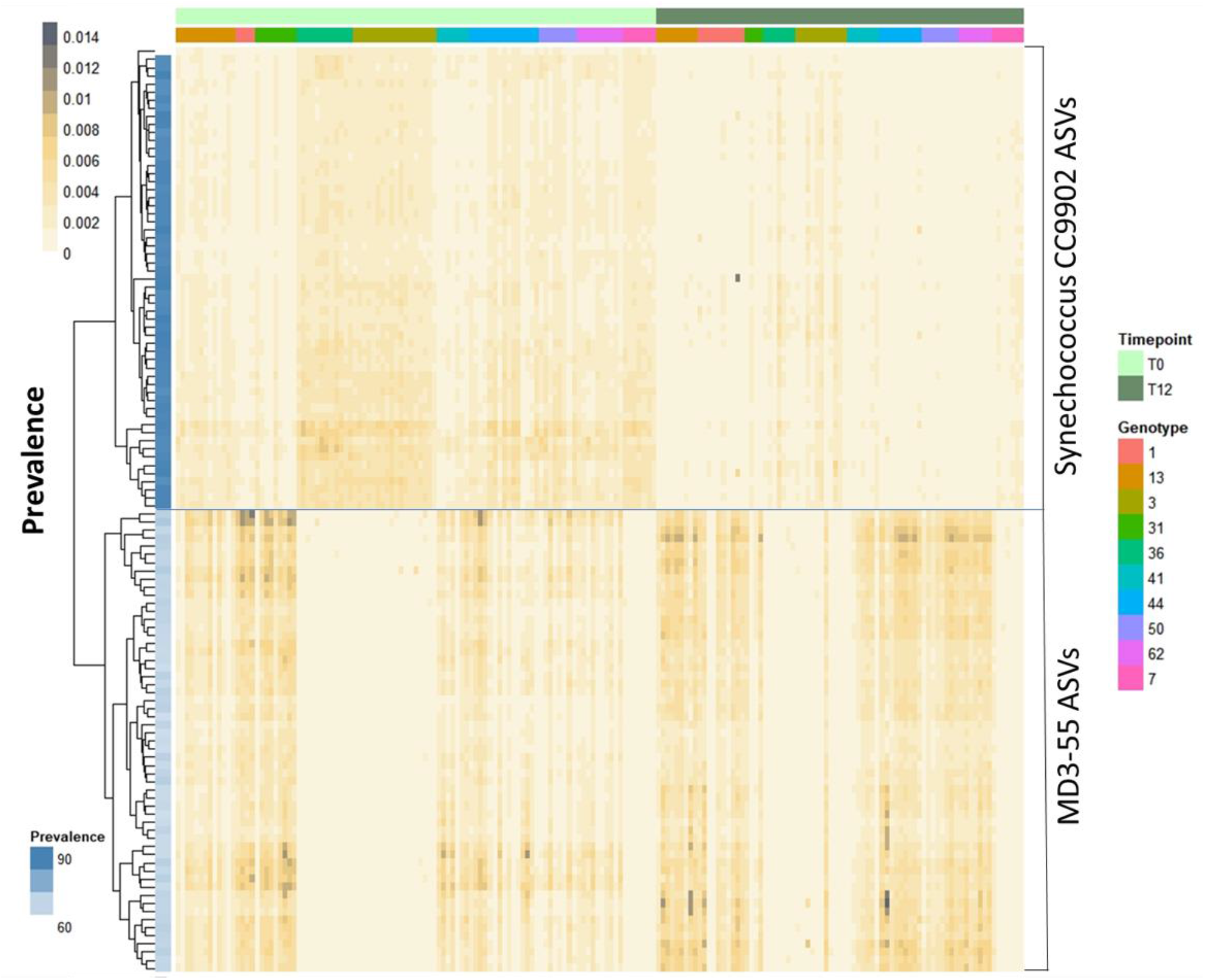
Heatmap displaying core taxa detected in samples at both timepoints, at a prevalence of >60%, and in frequencies of compositional relative abundance (light yellow=low to dark gray=high, gradient scale). Analysis was conducted with the microbiome package in R, using non-rarefied data. The two genera detected were also the most abundant in the dataset, MD3-55 and *Synechococcus* CC9902.

### ASVs of the ubiquitous intracellular symbiont, MD3-55 (sensu *A. rohweri*), increased one-year post-outplant

Phylogenetic classification placed our MD3-55 sequences alongside Rickettsiales symbiont, Candidatus *A. rohweri*, that was previously shown to affiliate with *A. cervicornis* hosts (Klinges et al. 2019; Baker et al. 2021), 99% of the time in 1000 bootstrap replicates (Fig. S4). The version of the SILVA database (version 138) which was used to assign taxonomy to our dataset has not updated the taxonomy of the Fokiniaceae family (genus MD3-55) to the newly proposed Candidatus *A. rohweri* (Klinges et al. 2019), therefore we will refer to these Rickettsiales as MD3-55.

While the pattern of low and high MD3-55 hosting *A. cervicornis* genotypes remained largely consistent over time, MD3-55 increased in abundance on average one year post transplantation (T_12_, Fig. 3a). This pattern was particularly evident for genotypes G13, G44, G50 and G62. Additionally, MD3-55 16S rRNA gene sequences diversified after 12 months, irrespective of outplant site, increasing from 70 distinct ASVs in T_0_ to 124 distinct ASVs in T_12_ samples (Fig. 3b). Of these, 57 variants of the MD3-55 population were observed at both timepoints (T_0_ and T_12_), while 13 variants disappeared, and 67 new variants were detected after 12 months (Fig. 3c).

**Figure 3.**
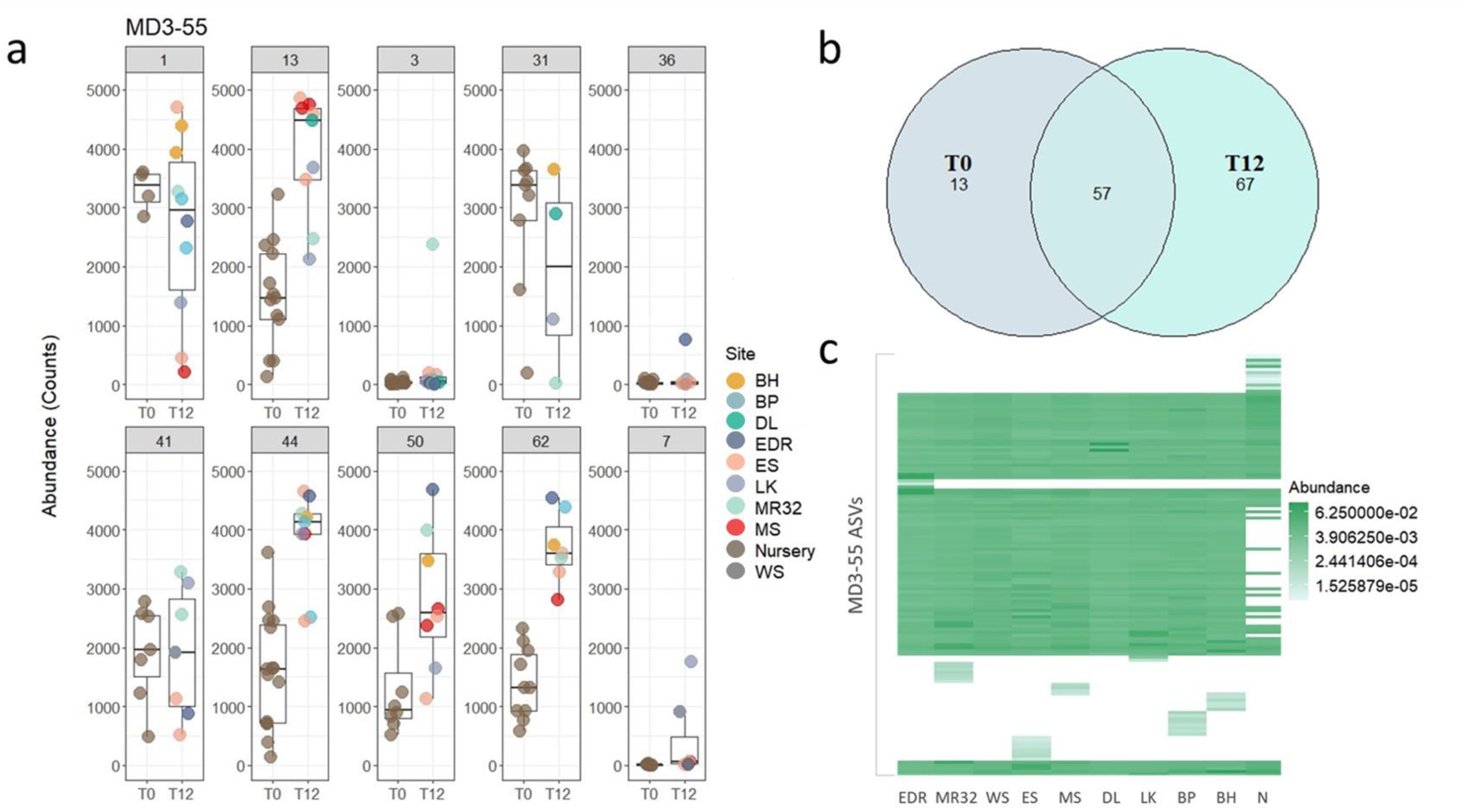
MD3-55 ASV distribution in the epibiomes of *A. cervicornis* genotypes. (a) Raw counts of MD3-55 in rarefied data, subset by genotypes. The x-axis denotes timepoints (T_0_ and T_12_) and transplant sites (BH: Bahia Honda, BP: Big Pine, DL: Dave’s Ledge, EDR: Eastern Dry Rocks, ES: Eastern Sambo, LK: Looe Key, MR32: Marker 32, MS: Maryland Shoals, N: Nursery, WS: Western Sambo) are differentiated by color. (b) Venn diagram of distinct and shared MD3-55 ASVs at T_0_ (nursery) and T_12_ (all transplant field sites, one year later). ASVs diversified following transplantation (67 distinct and 57 shared with the nursery samples), but 13 ASVs unique to the nursery were not detected at the transplant sites. (c) Heatmap of MD3-55 ASVs by transplant site (arranged from west to east) using non-metric multidimensional scaling and Bray-Curtis dissimilarity index. Each row represents a distinct MD3-55 ASV. Abundance is denoted by intensity of color, ranging from no abundance= white to high abundance= dark green.

To assess whether this increase in MD3-55 abundance was a real biological signal or an artifact of differential host tissue contamination, we tabulated the abundance of mitochondrial amplicons after pruning known protists from the dataset. Mitochondrial reads were higher in samples originating from the nursery than they were in the T_12_ field samples (Fig. S5) yet the opposite pattern was observed for MD3-55 (Fig. 3a).

## Discussion

Microbial communities are spatially organized between coral compartments and highly diverse (Sweet et al. 2011; Apprill et al. 2016; Hernandez-Agreda et al. 2017). Host-genotype specificity of the microbiome has been recently explored in acroporids but is still under-researched. Previous work on the tissue and mucus microbiome of various *Acropora* spp. has shown persistent host-genotype differences (Chu and Vollmer 2016; Glasl et al. 2019; Rosales et al. 2019). Although the epibiome of distinct *A. cervicornis* genotypes has been recently characterized (Miller et al. 2020), we show that genotype-specific differences in the mucus microbiome of *A. cervicornis* are largely maintained across space and time. The most abundant bacterial associates of the *A. cervicornis* microbial epibiotic community, both in the common garden nursery and following transplantation to nine distinct field sites, were members of the Fokiniaceae (MD3-55) and Synechococcaceae (*Synechococcus*). All other taxa (Fig. S1) were present in abundances of ≤10%, and their taxonomic composition did not seem to alter in transplants (Fig. S1), yet microbial signatures characteristic of low hosting MD3-55 genotypes remained (Fig.1b). Finally, MD3-55 symbionts were detected in the epibiome and increased over time in genotypes with initial high abundances.

### *A. cervicornis* epibiome retained genotype-specific signatures following transplantation

Site-specific differences in coral microbiomes are well documented (Rohwer et al. 2002; Guppy and Bythell 2006), and previous studies suggest reasonable flexibility of the acroporid microbiome under environmental change (Bourne et al. 2008; Grottoli et al. 2018). Coral mucus is susceptible to environmental effects (Li et al. 2015; Pollock et al. 2018; Marchioro et al. 2020), therefore we expected *A.cervicornis* epibiomes to be modulated by environmental parameters or geographic location and to highly resemble the seawater microbiome (Littman et al. 2009; Leite et al. 2018; Epstein et al. 2019). However, neither richness nor the microbiome composition at the phylum level were significantly altered in *A. cervicornis* genotypes following transplantation to different environments (Table S4, Fig. S3). (Marchioro et al. 2020) found that environmental parameters explained less of the variation in mucus microbiomes (10%) than that of microbial communities in the surrounding seawater (explained 32% of variation). Similarly, (Guppy and Bythell 2006) did not find strong correlations between environmental variables and bacterial structure of the mucus which led them to conclude that environmental influence was modulated by host intraspecific differences. Taken together, these results suggest host genotype may be a determining factor in structuring the epibiome of healthy corals, and that the environment may play a lesser role on mucus microbiomes than previously assumed.

Despite the strong signal of genotype, environmental influence on the epibiome cannot be completely ruled out, as different clustering patterns were observed for genotypes sampled in the nursery versus samples obtained from reefs, regardless of transplant site (Table S5, Fig. 1a). An increase in ASV richness was also observed in some genotypes after one year (Table S4), although the overall taxonomic composition remained the same. This pattern could be reflective of unique environmental differences in the nursery location, as it is a sand bottom habitat, but we did not resample genotypes in the nursery at T_12_ to test this hypothesis. Divergent clustering of the nursery-derived samples may also be related to climate variation between years as nursery and field samples were collected one year apart (April 2018 vs. April 2019). Transplants were visually monitored every three months for one year and no bleaching or disease was observed. Healthy coral tissue microbiomes can remain stable through time in coral species (Dunphy et al. 2019). The stability of the coral epibiome is unknown, but it is also possible that transplantation resulted in some level of dysbiosis leading to divergent microbiome composition regardless of the ultimate destination. Further research should integrate seasonal sampling in a time-series framework (2+ years) at nursery and field locations to disentangle spatial from temporal differences in coral and seawater microbiome variation and stability.

Additionally, the influence of microbial associations in near-coral seawater (seawater located within 5cm of corals) may play an important role in maintaining healthy, stable coral epibiomes through time in addition to influencing the microbiome of the surrounding seawater (Shashar et al. 1996; Weber et al. 2019). Weber et al. (2019) demonstrated that the coral ecosphere contains a taxonomically distinct, species-specific microbiome compared to that of seawater >1 m away from the reef. Our mucus samples were taken with syringes from the surface of the coral and most likely included near-coral seawater. FSW (nursery, BP and BH) controls resembled the microbial structure of low MD3-55 hosting genotypes (Fig S1), however, microbial communities were significantly different (Table S5), with the seawater containing more Bacteroidetes and Actinobacteria than coral mucus. It is also possible that the microbial structure of our FSW samples was influenced by the corals. Mucus shedding is a natural phenomenon in corals (Brown and Bythell 2005), and can be deployed during stressed conditions like increased UV exposure (Teai et al. 1998) and pathogen regulation (Ritchie 2006) but has also been linked to holobiont health via regulation of microbial communities (Glasl et al. 2016). It is possible that exuded dissolved organic matter from the mucus was present in the seawater (Silveira et al. 2017; Weber et al. 2019) and influenced the microbial structure of our FSW samples since seawater was collected > 1 meter of the focal corals. This may explain why the only distinguishing microbial signature between corals and seawater the presence/absence of MD3-55 (Fig.1b) was. Lastly, microbial beta-diversity in the seawater did not significantly differ among sites (Table S5) despite the one-year difference in sampling time between the nursery and BH/BP sites. The sites are ~8km apart and the nursery is closest to LK (Table S1). The similarities in seawater microbiomes indicate that variation in physicochemical conditions did not significantly affect the local microbiota between sites. Additionally, we did not observe drastic variation in the microbial structure of *A. cervicornis* epibiomes by site (other than differences in the presence/absence of specific MD3-55 ASVs), only genotype effects (Fig. 1a). In order to assess the extent of environmental effects on the mucus microbiome of *A. cervicornis* genotypes, and vice-versa, follow-up studies should monitor physicochemical parameters during sampling and sample seawater at least one meter away from the reef.

### MD3-55 in the epibiome

The putative bacterial symbiont, MD3-55, is ubiquitously present in *A. cervicornis* (Casas et al. 2004; Miller et al. 2014; Gignoux-Wolfsohn and Vollmer 2015). No significant differences were observed among genotypes when MD3-55 ASVs were removed from the samples (Table S5). These findings point to MD3-55 as the driving factor distinguishing genotype-specific epibiomes, similar to those signatures reported for A*. cervicornis* tissue samples (Klinges et al. 2020; Miller et al. 2020; Baker et al. 2021). MD3-55 were previously visualized (Gignoux-Wolfsohn et al. 2020), and are known to lack basic metabolic pathways, strongly supporting an intracellular lifestyle (Klinges et al. 2019). However, the localization of MD3-55 in *A. cervicornis* remains elusive. Baker et al. (2021) hypothesized *A. cervicornis* may horizontally transmit MD3-55 to gametes and juveniles via mucocytes. A histological approach supports this hypothesis as Rickettsiales-like organisms (RLO) were visualized in the mucocytes of *A. cervicornis* (Miller et al. 2014; Gignoux-Wolfsohn et al. 2020). Our study taxonomically identified MD3-55 in the epibiome, suggesting that those previously identified RLOs located in and near the mucocytes may be MD3-55 which would indicate that these organisms are not exclusively intracellular.

Although we observed MD3-55 in surface mucus samples, they could have derived from host tissue contamination. To address this issue, we assessed mitochondrial reads to evaluate whether host tissue contamination increased across sampling time-points. Mitochondrial reads did not increase in our T_12_ data and were lower in the T_12_ sample set than the T_0_ dataset (Fig. S5), whereas MD3-55 increased in T_12_ in G13, G44, G50 and G62 (Fig. 3a), which argues against host tissue contamination. This suggests that MD3-55 may display partial extracellular inclinations, perhaps to facilitate horizontal transmission. Additionally, MD3-55 is not host restricted and has been detected in sponges, kelp, ctenophores, and marine sediments. Investigating the abundance and localization of MD3-55 in non-coral hosts residing in the same habitats as acroporids, and other environmental reservoirs, may be valuable in resolving the general ecology of MD3-55.

### MD3-55 ASVs following transplantation in high MD3-55 hosting genotypes

Recent work has shown that MD3-55 is highly abundant (Klinges et al. 2020; Baker et al. 2021) in genotypes of *A. cervicornis*, previously determined to be susceptible to white-band disease after a bleaching event caused by a temperature stress (Muller et al. 2018). White-band is a devastating, host-specific disease in *A. cervicornis* and *A. palmata* with an unknown etiological origin, although the putative pathogen is likely bacterial (Casas et al. 2004; Kline and Vollmer 2011; Sweet et al. 2014; Gignoux-Wolfsohn and Vollmer 2015).

Here, we observed an increase in MD3-55 in several high MD3-55 hosting genotypes following transplantation to novel field sites. While there was some indication that MD3-55 increased in some ramets of low MD3-55 hosting genotypes, these genotypes largely maintained a persistent signature of low MD3-55 abundance (Fig. 3a). Recently, Baker et al. (2021) indicated *A. rohweri* populations in *A. cervicornis* were abundantly high and had greater *in situ* replication rates in Florida (USA) compared to those from Belize and the U.S. Virgin Islands. We also observed several nursery MD3-55 ASVs disappear following transplantation, while others proliferated. Proliferation of particular MD3-55 ASVs also appeared to be site specific (Fig. 3c) suggesting positive selection in certain locations, as previously observed in Florida (higher abundances) and Caribbean (lower abundances) populations of MD3-55 in *A. cervicornis* and *A. prolifera* (Baker et al. 2021). Higher abundances were attributed to environmental factors, like higher nutrient stress, which aligns with our observations of host-specific abundance patterns of MD3-55 out in the field. However, we did not quantify environmental parameters like dissolved nutrients (nitrogen, phosphorus) or particulate organic carbon, therefore we cannot conclusively link high MD3-55 abundance patterns in our data to increased nutrient availability at specific field sites.

We identified 137 MD3-55 ASVs, (Fig 3b) in our ten *A. cervicornis* genotypes whereas (Miller et al. 2020) identified 11 MD3-55 ASVs in three genotypes (different from those in our dataset) using the same 16S rRNA amplifying primers and sequencing protocol. In the nursery, we observed 70 initial ASVs and a diversification of 67 novel ASVs in the field sites after one year (Fig. 3b, c). Baker et al. (2021) showed greater positive selection in genes associated with ribosomal assembly in MD3-55 strains from Florida, signaling possible speciation across locations. These findings, along with our study, suggest rapid evolution and diversification in MD3-55 strains may be happening in Florida field sites. Novel infections in low MD3-55 hosting genotypes were detected in three different sites, but only in 1 ramet each in G3, G36 and 2 ramets in G7 (Fig. 3a). Local acquisition via the surrounding seawater or sediments may be a possibility, but we observed none to very low abundances of MD3-55 in FSW samples of the nursery and field sites BH and BP (Fig. 1b).

### Persistence of cyanobacteria in the epibiome

The most abundant taxa, MD3-55 and *Synechococcus* CC9902 (Fig. 1b), in *A. cervicornis* epibiomes were also the most stable, with MD3-55 ASVs present in >60% of the samples (consistent with the observation of high and low MD3-55 hosting genotypes) and *Synechococcus* ASVs present in ~90% (consistent in all genotypes) of the samples (Fig.2). This latter finding is consistent with a recent report on *A. tenuis* and *A. millepora* microbiomes where *Synechococcus* was documented at higher abundances in the mucus than in the tissue and surrounding seawater (Marchioro et al. 2020).

While the ecology and functional role of MD3-55 in *A. cervicornis* may involve parasitism (Klinges et al. 2019, 2020), it is not well understood. In contrast, the associations between cyanobacteria and corals are putatively related to nitrogen fixation (Lesser 2004; Lesser et al. 2007; Lema et al. 2012). Cyanobacteria can establish partnerships with various organisms, such as other prokaryotes, microbial eukaryotes and metazoans (Mutalipassi et al. 2021). In most cases the bulk of the exchanged services involve biologically useful nitrogen (Foster and O’Mullan 2008). For example, δ15N stable isotope data suggests algal symbionts preferentially use nitrogen fixed by cyanobacteria, including *Synechococcus*, in colonies of the coral, *Montastraea cavernosa* (Lesser et al. 2007). It is unknown if algal symbionts of *A.cervicornis* share a similar nitrogen acquisition strategy but cyanobacteria nitrogen-fixers are ubiquitous in the tissue and mucus of acroporids from the Great Barrier Reef and Caribbean (Kvennefors and Roff 2009; Lema et al. 2012; Marchioro et al. 2020; Miller et al. 2020). Other cyanobiont-mediated services have also been identified in healthy corals, like the exchange of photoprotective compounds in *Montastraea cavernosa* (Lesser 2004; Lesser et al. 2007). Although some cyanobacteria have been associated as precursors to black-band disease (BBD) (Frias-Lopez et al. 2003), a bacterial mat that kills and removes healthy tissue and beneficial bacterial associates from corals (Richardson 1996; Gantar et al. 2011), *Synechococcus* species are not linked to BBD (Klaus et al. 2011; Buerger et al. 2016). Given that all of our coral were visibly healthy at the time of sampling and *Synechococcus* were also present in FSW site samples, the association between *A. cervicornis* and *Synechococcus* is likely mutualistic or commensal. To explore this, future work should investigate host-specific distributions of cyanobacteria in *A. cervicornis* and explore the role of cyanobiont-mediated nitrogen in maintenance of the cnidarian-algal symbiosis.

In summary, understanding the influence of host specificity and the environment on the maintenance of acroporid epibiomes is pivotal if microbial markers are to be used in reef restoration (Parkinson et al. 2020). Prior work in *A. tenuis* suggests that significant genotype variability may limit the use of microbiome surveys as microbial indicators of coral colony health (Glasl et al. 2019). Here, we also observe significant and persistent variation in the composition of the mucus microbiome among genotypes of *A. cervicornis*, but this does not necessarily preclude the potential utility of microbial indicators. Disease-resistant and disease-susceptible *A. cervicornis* genotypes retain distinct tissue microbiome signatures, with MD3-55 as the primary differentiating “biomarker” (Klinges et al. 2020). Although MD3-55 was only recently detected in coral epibiomes (Miller et al. 2020), we show that the presence and abundance of MD3-55 in *A. cervicornis* genotypes can also be reliably detected in the epibiome throughout time. Moreover, high and low infection types are retained through different environmental exposures over time. This result has particular utility for reef restoration applications, as non-invasive sampling of the mucus and surface microbiome of threatened *A. cervicornis* can potentially inform on the disease-susceptibility or disease-resistance of restored populations in natural environments.

## Acknowledgements

This study was funded by National Oceanic and Atmospheric Administration Coral Reef Conservation Program grant NA17NOS4820084 and National Science Foundation Graduate Research Fellowship Program grant award DGE-1418060. We would like to thank Y. Zhang for help with collections. A. Clark kindly provided access to a vacuum pump. Thanks to the Florida Keys National Marine Sanctuary for authorizing this work under FKNMS permits 2015-163-A1 and 2018-035.

## Contributions

CDK and CJK conceived and designed the field experiment and obtained funding. EGA and CJK performed extractions. EGA completed library preparation, all bioinformatic and statistical analyses and wrote the first draft of the manuscript. All authors contributed to sample collection and revisions.

## Data Accessibility

Scripts for data analysis used in this project are available at https://github.com/symbiotic-em/acer_epi_final. Demultiplexed sequences are available at the National Center for Biotechnology Information (NCBI) Sequence Read Archives (SRA) under accession code: PRJNA630333.

## References

Andrews S. (2010) FastQC: A Quality Control Tool for High Throughput Sequence Data. http://www.bioinformatics.babraham.ac.uk/projects/fastqc/

Apprill A, Weber LG, Santoro AE (2016) Distinguishing between Microbial Habitats Unravels Ecological Complexity in Coral Microbiomes. mSystems 1:

Baker LJ, Reich HG, Kitchen SA, Grace Klinges J, Koch HR, Baums IB, Muller E, Thurber RV (2021) The coral symbiont Candidatus Aquarickettsia is variably abundant in threatened Caribbean acroporids and transmitted horizontally.

Blackall LL, Wilson B, van Oppen MJH (2015) Coral-the world’s most diverse symbiotic ecosystem. Mol Ecol 24:5330–5347

Bourne DG, Morrow KM, Webster NS (2016) Insights into the Coral Microbiome: Underpinning the Health and Resilience of Reef Ecosystems. Annu Rev Microbiol 70:317–340

Bourne D, Iida Y, Uthicke S, Smith-Keune C (2008) Changes in coral-associated microbial communities during a bleaching event. The ISME Journal 2:350–363

Brown BE, Bythell JC (2005) Perspectives on mucus secretion in reef corals. Marine Ecology Progress Series 296:291–309

Buerger P, Alvarez-Roa C, Weynberg KD, Baekelandt S, van Oppen MJH (2016) Genetic, morphological and growth characterisation of a newRoseofilumstrain (Oscillatoriales, Cyanobacteria) associated with coral black band disease. PeerJ 4:e2110

Callahan BJ, McMurdie PJ, Rosen MJ, Han AW, Johnson AJA, Holmes SP (2016) DADA2: High-resolution sample inference from Illumina amplicon data. Nat Methods 13:581–583

Caporaso JG, Lauber CL, Walters WA, Berg-Lyons D, Lozupone CA, Turnbaugh PJ, Fierer N, Knight R (2011)Global patterns of 16S rRNA diversity at a depth of millions of sequences per sample. Proc Natl Acad Sci U S A 108 Suppl 1:4516–4522

Casas V, Kline DI, Wegley L, Yu Y, Breitbart M, Rohwer F (2004) Widespread association of a Rickettsiales-like bacterium with reef-building corals. Environ Microbiol 6:1137–1148

Chu ND, Vollmer SV (2016) Caribbean corals house shared and host-specific microbial symbionts over time and space. Environ Microbiol Rep 8:493–500

Coffroth MA (1990) Mucous sheet formation on poritid corals: An evaluation of coral mucus as a nutrient source on reefs. Marine Biology 105:39–49

Crossland CJ (1987) In situ release of mucus and DOC-lipid from the corals Acropora variabilis and Stylophora pistillata in different light regimes. Coral Reefs 6:35–42

Dunphy CM, Gouhier TC, Chu ND, Vollmer SV (2019) Structure and stability of the coral microbiome in space and time. Sci Rep 9:6785

Edgar RC (2004) MUSCLE: a multiple sequence alignment method with reduced time and space complexity. BMC Bioinformatics 5:113

Epstein HE, Smith HA, Cantin NE, Mocellin VJL, Torda G, van Oppen MJH (2019) Temporal Variation in the Microbiome of Acropora Coral Species Does Not Reflect Seasonality. Front Microbiol 10:

Foster RA, O’Mullan GD (2008) Nitrogen-Fixing and Nitrifying Symbioses in the Marine Environment. Nitrogen in the Marine Environment 1197–1218

Frias-Lopez J, Bonheyo GT, Jin Q, Fouke BW (2003) Cyanobacteria Associated with Coral Black Band Disease in Caribbean and Indo-Pacific Reefs. Applied and Environmental Microbiology 69:2409–2413

Gantar M, Kaczmarsky LT, Stanić D, Miller AW, Richardson LL (2011) Antibacterial activity of marine and black band disease cyanobacteria against coral-associated bacteria. Mar Drugs 9:2089–2105

Garren M, Azam F (2010) New Method for Counting Bacteria Associated with Coral Mucus. Applied and Environmental Microbiology 76:6128–6133

Gignoux-Wolfsohn SA, Precht WF, Peters EC, Gintert BE, Kaufman LS (2020) Ecology, histopathology, and microbial ecology of a white-band disease outbreak in the threatened staghorn coral Acropora cervicornis. Dis Aquat Organ 137:217–237

Gignoux-Wolfsohn SA, Vollmer SV (2015) Identification of Candidate Coral Pathogens on White Band Disease-Infected Staghorn Coral. PLoS One 10:e0134416

Glasl B, Herndl GJ, Frade PR (2016) The microbiome of coral surface mucus has a key role in mediating holobiont health and survival upon disturbance. ISME J 10:2280–2292

Glasl B, Smith CE, Bourne DG, Webster NS (2019) Disentangling the effect of host-genotype and environment on the microbiome of the coral. PeerJ 7:e6377

Gloor GB, Macklaim JM, Pawlowsky-Glahn V, Egozcue JJ (2017) Microbiome Datasets Are Compositional: And This Is Not Optional. Front Microbiol 8:2224

Grottoli AG, Dalcin Martins P, Wilkins MJ, Johnston MD, Warner ME, Cai W-J, Melman TF, Hoadley KD, Pettay DT, Levas S, Schoepf V (2018) Coral physiology and microbiome dynamics under combined warming and ocean acidification. PLoS One 13:e0191156

Guppy R, Bythell JC (2006) Environmental effects on bacterial diversity in the surface mucus layer of the reef coral Montastraea faveolata. Marine Ecology Progress Series 328:133–142

Hernandez-Agreda A, Gates RD, Ainsworth TD (2017) Defining the Core Microbiome in Corals’ Microbial Soup. Trends in Microbiology 25:125–140

Kalyaanamoorthy S, Minh BQ, Wong TKF, von Haeseler A, Jermiin LS (2017) ModelFinder: fast model selection for accurate phylogenetic estimates. Nature Methods 14:587–589

Kauffmann A, Gentleman R, Huber W (2009) arrayQualityMetrics--a bioconductor package for quality assessment of microarray data. Bioinformatics 25:415–416

Klaus JS, Janse I, Fouke BW (2011) Coral Black Band Disease Microbial Communities and Genotypic Variability of the Dominant Cyanobacteria (CD1C11). Bulletin of Marine Science 87:795–821

Kline DI, Vollmer SV (2011) White Band Disease (type I) of endangered caribbean acroporid corals is caused by pathogenic bacteria. Sci Rep 1:7

Klinges G, Maher RL, Vega Thurber RL, Muller EM (2020) Parasitic “Candidatus Aquarickettsia rohweri” is a marker of disease susceptibility in Acropora cervicornis but is lost during thermal stress. Environ Microbiol 22:5341–5355

Klinges JG, Rosales SM, McMinds R, Shaver EC, Shantz AA, Peters EC, Eitel M, Wörheide G, Sharp KH, Burkepile DE, Silliman BR, Vega Thurber RL (2019) Phylogenetic, genomic, and biogeographic characterization of a novel and ubiquitous marine invertebrate-associated Rickettsiales parasite, Candidatus Aquarickettsia rohweri, gen. nov., sp. nov. ISME J 13:2938–2953

Kolde R, Franzosa EA, Rahnavard G, Hall AB, Vlamakis H, Stevens C, Daly MJ, Xavier RJ, Huttenhower C (2018)Host genetic variation and its microbiome interactions within the Human Microbiome Project. Genome Med 10:6

Kooperman N, Ben-Dov E, Kramarsky-Winter E, Barak Z, Kushmaro A (2007) Coral mucus-associated bacterial communities from natural and aquarium environments. FEMS Microbiol Lett 276:106–113

Krediet CJ, Ritchie KB, Paul VJ, Teplitski M (2013) Coral-associated micro-organisms and their roles in promoting coral health and thwarting diseases. Proc Biol Sci 280:20122328

Kvennefors ECE, Roff G (2009) Evidence of cyanobacteria-like endosymbionts in Acroporid corals from the Great Barrier Reef. Coral Reefs 28:547–547

Leite DCA, Salles JF, Calderon EN, Castro CB, Bianchini A, Marques JA, van Elsas JD, Peixoto RS (2018) Coral Bacterial-Core Abundance and Network Complexity as Proxies for Anthropogenic Pollution. Front Microbiol 9:833

Lema KA, Willis BL, Bourne DG (2012) Corals form characteristic associations with symbiotic nitrogen-fixing bacteria. Appl Environ Microbiol 78:3136–3144

Lahti L, Shetty S (2012-2019) microbiome R package. http://microbiome.github.io

Lesser MP (2004) Discovery of Symbiotic Nitrogen-Fixing Cyanobacteria in Corals. Science 305:997–1000

Lesser MP, Falcón LI, Rodríguez-Román A, Enríquez S, Hoegh-Guldberg O, Iglesias-Prieto R (2007) Nitrogen fixation by symbiotic cyanobacteria provides a source of nitrogen for the scleractinian coral Montastraea cavernosa. Marine Ecology Progress Series 346:143–152

Li J, Chen Q, Long L-J, Dong J-D, Yang J, Zhang S (2015) Bacterial dynamics within the mucus, tissue and skeleton of the coral Porites lutea during different seasons. Scientific Reports 4:

Littman RA, Willis BL, Pfeffer C, Bourne DG (2009) Diversities of coral-associated bacteria differ with location, but not species, for three acroporid corals on the Great Barrier Reef. FEMS Microbiol Ecol 68:152–163

Love MI, Huber W, Anders S (2014) Moderated estimation of fold change and dispersion for RNA-seq data with DESeq2. Genome Biol 15:550

Lynch JB, Hsiao EY (2019) Microbiomes as sources of emergent host phenotypes. Science 365:1405–1409

Marchioro GM, Glasl B, Engelen AH, Serrão EA, Bourne DG, Webster NS, Frade PR (2020) Microbiome dynamics in the tissue and mucus of acroporid corals differ in relation to host and environmental parameters. PeerJ 8:e9644

McDevitt-Irwin JM, Baum JK, Garren M, Vega Thurber RL (2017) Responses of Coral-Associated Bacterial Communities to Local and Global Stressors. Frontiers in Marine Science 4:

McMurdie PJ, Holmes S (2013) phyloseq: an R package for reproducible interactive analysis and graphics of microbiome census data. PLoS One 8:e61217

Meikle P, Richards GN, Yellowlees D (1988) Structural investigations on the mucus from six species of coral. Marine Biology 99:187–193

Miller MW, Lohr KE, Cameron CM, Williams DE, Peters EC (2014) Disease dynamics and potential mitigation among restored and wild staghorn coral, Acropora cervicornis. PeerJ 2:e541

Miller N, Maneval P, Manfrino C, Frazer TK, Meyer JL (2020) Spatial distribution of microbial communities among colonies and genotypes in nursery-reared. PeerJ 8:e9635

Minh BQ, Schmidt HA, Chernomor O, Schrempf D, Woodhams MD, von Haeseler A, Lanfear R (2020) IQ-TREE 2: New Models and Efficient Methods for Phylogenetic Inference in the Genomic Era. Mol Biol Evol 37:1530–1534

Muller EM, Bartels E, Baums IB (2018) Bleaching causes loss of disease resistance within the threatened coral species. Elife 7:

Mutalipassi M, Riccio G, Mazzella V, Galasso C, Somma E, Chiarore A, de Pascale D, Zupo V (2021) Symbioses of Cyanobacteria in Marine Environments: Ecological Insights and Biotechnological Perspectives. Mar Drugs 19:

Oksanen J, Blanchet G, Friendly M, Kindt R, Legendre P, McGlinn D, Minchin PR, O’Hara RB, Simpson GL, Solymos P, Stevens MHH, Szoecs E, Wagner H (2019) vegan: Community Ecology Package. R package version 2.5-6. https://CRAN.R-project.org/package=vegan

van Oppen MJH, Blackall LL (2019) Coral microbiome dynamics, functions and design in a changing world. Nat Rev Microbiol 17:557–567

Parada AE, Needham DM, Fuhrman JA (2016) Every base matters: assessing small subunit rRNA primers for marine microbiomes with mock communities, time series and global field samples. Environ Microbiol 18:1403–1414

Parkinson JE, Baker AC, Baums IB, Davies SW, Grottoli AG, Kitchen SA, Matz MV, Miller MW, Shantz AA, Kenkel CD (2020) Molecular tools for coral reef restoration: Beyond biomarker discovery. Conservation Letters 13:e12687

Peixoto RS, Rosado PM, Leite DC de A, Rosado AS, Bourne DG (2017) Beneficial Microorganisms for Corals(BMC): Proposed Mechanisms for Coral Health and Resilience. Front Microbiol 8:341

Pollock FJ, Joseph Pollock F, Lamb JB, Field SN, Heron SF, Schaffelke B, Shedrawi G, Bourne DG, Willis BL(2014) Sediment and Turbidity Associated with Offshore Dredging Increase Coral Disease Prevalence on Nearby Reefs. PLoS ONE 9:e102498

Pollock FJ, McMinds R, Smith S, Bourne DG, Willis BL, Medina M, Thurber RV, Zaneveld JR (2018) Coral-associated bacteria demonstrate phylosymbiosis and cophylogeny. Nat Commun 9:4921

R Core Team (2020) R: A language and environment for statistical computing. R Foundation for Statistical Computing, Vienna, Austria. https://www.R-project.org

Richardson LL (1996) Horizontal and vertical migration patterns of Phorrnidium corallyticum and Beggiatoa spp.associated with black-band disease of corals. Microbial Ecology 32:

Ritchie KB (2006) Regulation of microbial populations by coral surface mucus and mucus-associated bacteria. Marine Ecology Progress Series 322:1–14

Rohwer F, Seguritan V, Azam F, Knowlton N (2002) Diversity and distribution of coral-associated bacteria. Marine Ecology Progress Series 243:1–10

Rosado PM, Leite DCA, Duarte GAS, Chaloub RM, Jospin G, Nunes da Rocha U, P Saraiva J, Dini-Andreote F, Eisen JA, Bourne DG, Peixoto RS (2019) Marine probiotics: increasing coral resistance to bleaching throughm icrobiome manipulation. ISME J 13:921–936

Rosales SM, Miller MW, Williams DE, Traylor-Knowles N, Young B, Serrano XM (2019) Microbiome differences in disease-resistant vs. susceptible Acropora corals subjected to disease challenge assays. Sci Rep 9:18279

Shashar N, Kinane S, Jokiel PL, Patterson MR (1996) Hydromechanical boundary layers over a coral reef. Journal of Experimental Marine Biology and Ecology 199:17–28

Silveira CB, Cavalcanti GS, Walter JM, Silva-Lima AW, Dinsdale EA, Bourne DG, Thompson CC, Thompson FL(2017) Microbial processes driving coral reef organic carbon flow. FEMS Microbiol Rev 41:575–595

Sutherland KP, Porter JW, Torres C (2004) Disease and immunity in Caribbean and Indo-Pacific zooxanthellate corals. Marine Ecology Progress Series 266:273–302

Sweet MJ, Bulling MT (2017) On the Importance of the Microbiome and Pathobiome in Coral Health and Disease. Frontiers in Marine Science 4:

Sweet MJ, Croquer A, Bythell JC (2011) Bacterial assemblages differ between compartments within the coral holobiont. Coral Reefs 30:39–52

Sweet MJ, Croquer A, Bythell JC (2014) Experimental antibiotic treatment identifies potential pathogens of white band disease in the endangered Caribbean coral Acropora cervicornis. Proc Biol Sci 281:20140094

Teai T, Drollet JH, Bianchini J-P, Cambon A, Martin PMV (1998) Occurrence of ultraviolet radiation-absorbing mycosporine-like amino acids in coral mucus and whole corals of French Polynesia. Marine and Freshwater Research 49:127

Thompson LR, The Earth Microbiome Project Consortium, Sanders JG, McDonald D, Amir A, Ladau J, Locey KJ, Prill RJ, Tripathi A, Gibbons SM, Ackermann G, Navas-Molina JA, Janssen S, Kopylova E, Vázquez-Baeza Y, González A, Morton JT, Mirarab S, Xu ZZ, Jiang L, Haroon MF, Kanbar J, Zhu Q, Song SJ, Kosciolek T, Bokulich NA, Lefler J, Brislawn CJ, Humphrey G, Owens SM, Hampton-Marcell J, Berg-Lyons D, McKenzie V, Fierer N, Fuhrman JA, Clauset A, Stevens RL, Shade A, Pollard KS, Goodwin KD, Jansson JK, Gilbert JA, Knight R (2017) A communal catalogue reveals Earth’s multiscale microbial diversity. Nature 551:457–463

Vogel KJ, Moran NA (2011) Effect of Host Genotype on Symbiont Titer in the Aphid-Buchnera Symbiosis. Insects 2:423–434

Wagner MR, Lundberg DS, Del Rio TG, Tringe SG, Dangl JL, Mitchell-Olds T (2016) Host genotype and age shape the leaf and root microbiomes of a wild perennial plant. Nat Commun 7:12151

Wang Q, Garrity GM, Tiedje JM, Cole JR (2007) Naive Bayesian classifier for rapid assignment of rRNA sequences into the new bacterial taxonomy. Appl Environ Microbiol 73:5261–5267

Weber L, Gonzalez-Díaz P, Armenteros M, Apprill A (2019) The coral ecosphere: A unique coral reef habitat that fosters coral-microbial interactions. Limnology and Oceanography 64:2373–2388

Weiss S, Xu ZZ, Peddada S, Amir A, Bittinger K, Gonzalez A, Lozupone C, Zaneveld JR, Vázquez-Baeza Y, Birmingham A, Hyde ER, Knight R (2017) Normalization and microbial differential abundance strategies depend upon data characteristics. Microbiome 5:27

Wickham H (2016) ggplot2: Elegant graphics for data analysis. Springer-Verlag New York

Wild C, Huettel M, Klueter A, Kremb SG, Rasheed MYM, Jørgensen BB (2004) Coral mucus functions as an energy carrier and particle trap in the reef ecosystem. Nature 428:66–70

Wright RM, Kenkel CD, Dunn CE, Shilling EN, Bay LK, Matz MV (2017) Intraspecific differences in molecular stress responses and coral pathobiome contribute to mortality under bacterial challenge in Acropora millepora. Sci Rep 7:2609

