## Supplemental Figures S1-S5, Tables S1-S6 for "Host-specific epibiomes of distinct *Acropora cervicornis* genotypes persist after field transplantation"

**SUPPLEMENTARY FILE**

Figures S1-S5

Tables S1-Table S6

**Host-specific epibiomes of distinct *Acropora cervicornis* genotypes persist after field transplantation**

Emily G. Aguirre*^1^, Wyatt C. Million^1^, Erich Bartels^2^, Cory J. Krediet^3^, Carly D. Kenkel^1^

^1^ Department of Biological Sciences, University of Southern California, 3616 Trousdale Parkway, Los Angeles, CA 90089, United States of America

^2^ Elizabeth Moore International Center for Coral Reef Research & Restoration, Mote Marine Laboratory, 24244 Overseas Hwy, Summerland Key, FL 33042, United States

^3^ Department of Marine Science, Eckerd College, 4200 54th Avenue South, St. Petersburg, FL 33711, United States of America

*Corresponding author:

Emily G Aguirre

University of Southern California

3616 Trousdale Parkway

Los Angeles, CA 90089, USA

**Figure S1.** Relative abundance of all taxa, by phylum, in *A. cervicornis* genotypes and background FSW (N: Nursery, BP: Big Pine Shoals, BH: Bahia Honda, LK: Looe Key). The predominant taxa are Proteobacteria, followed by cyanobacteria and actinobacteria. All other taxa were present at < 10% relative abundance.
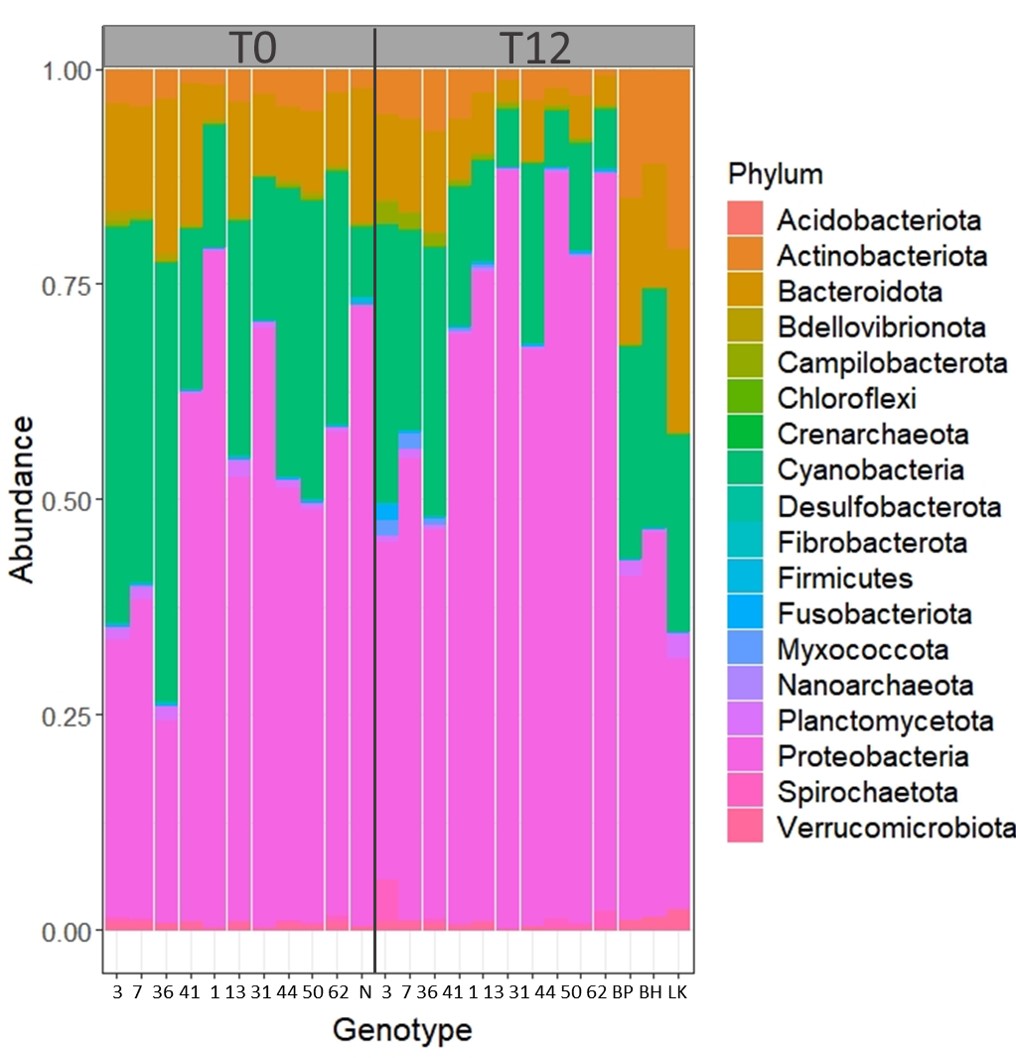

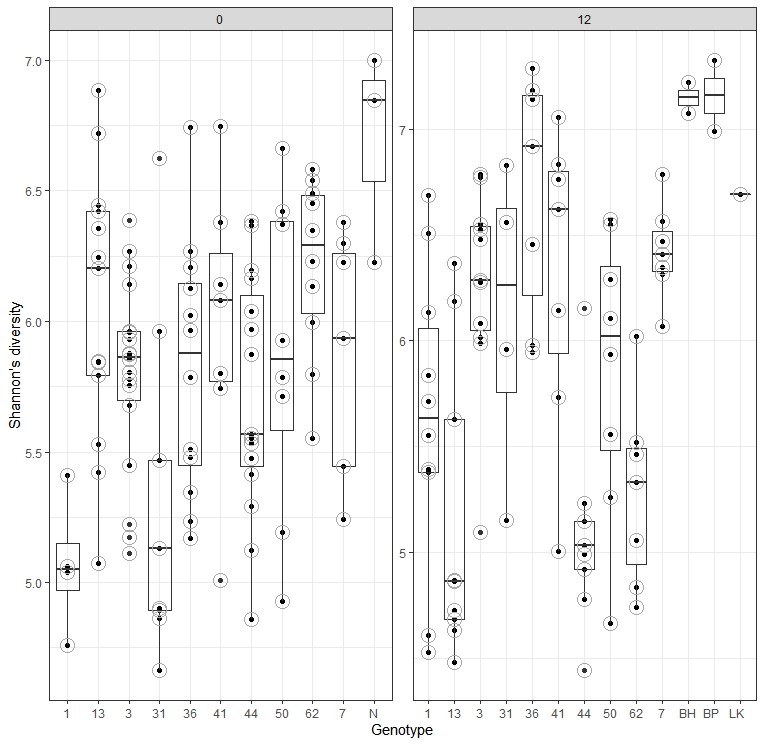

**Figure S2.** Alpha-diversity of the genotypes and background FSW (N: Nursery, BP: Big Pine Shoals, BH: Bahia Honda, LK: Looe Key) at the nursery (0) and transplant sites (12), measured using Shannon’s index.

**Figure S3.** Relative abundance of all taxa, by phylum, in samples sorted by host genotype and background FSW (BP: Big Pine Shoals, BH: Bahia Honda, LK: Looe Key), and by transplant sites at T_12_ only. Differences in relative abundance of phyla can be seen in the genotypes but is indistinguishable if samples are visualized only by site.
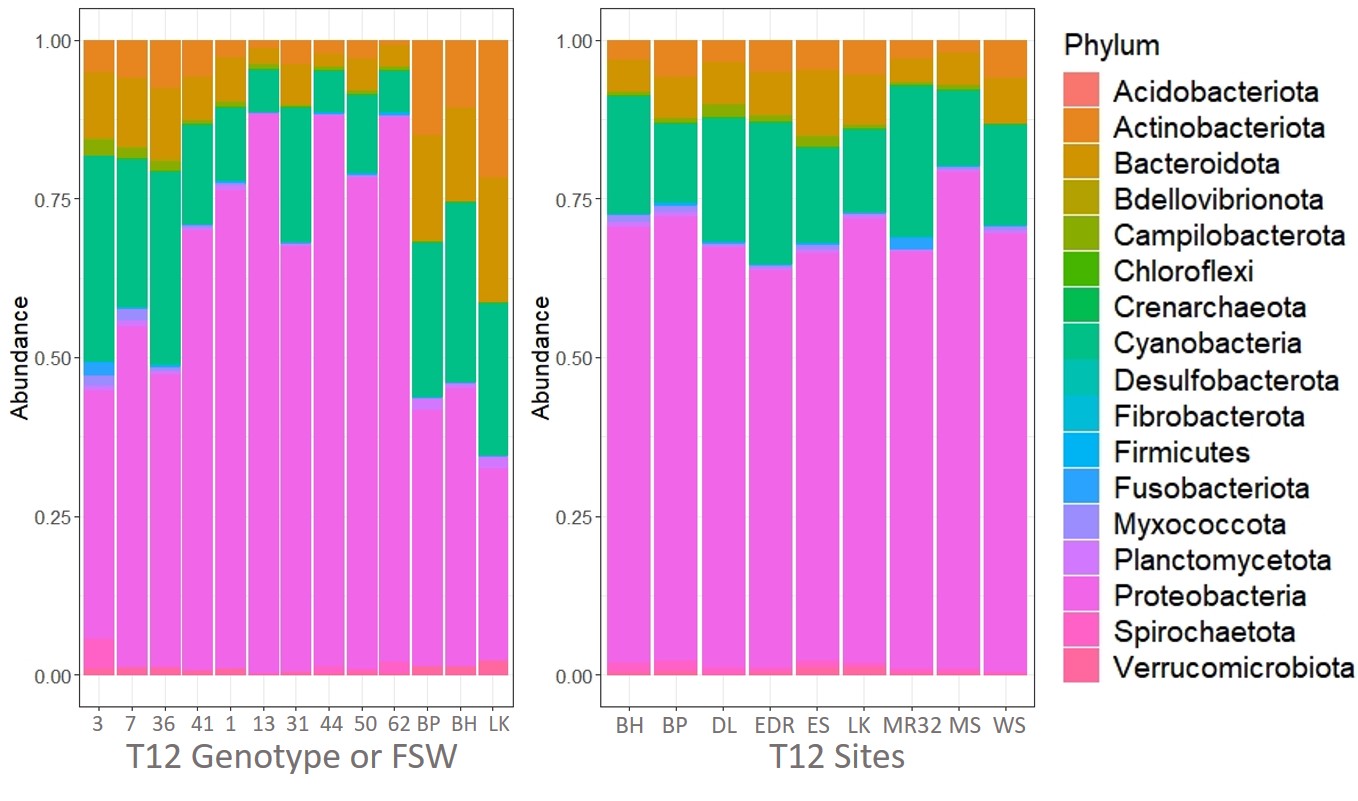

**Figure S4.** Phylogenetic classification of 16S rRNA MD3-55 SILVA-assigned sequences (red text and circles) with two published 16S rRNA sequences of coral-associated Fokiniaceae (“MD3.55 16S NCBI” and “*Aquarickettsia rohwer*i”) and other members of the Rickettsiales (12 families). *Caulobacter mirabilis* (Alphaproteobacteria), was chosen as the outgroup representative. Phylogenetic classification was conducted using IQTREE. Briefly, a maximum likelihood phylogeny with ultrafast bootstrap (1000 reps.) was applied and tree visualization was done on the Interactive Tree of Life website. Only one of our sequences (red ASV54) clustered with the MD3-55 NCBI sequence, whereas the remaining MD3-55 ASVs clustered with the published *A. rohweri* 16S rRNA sequence.
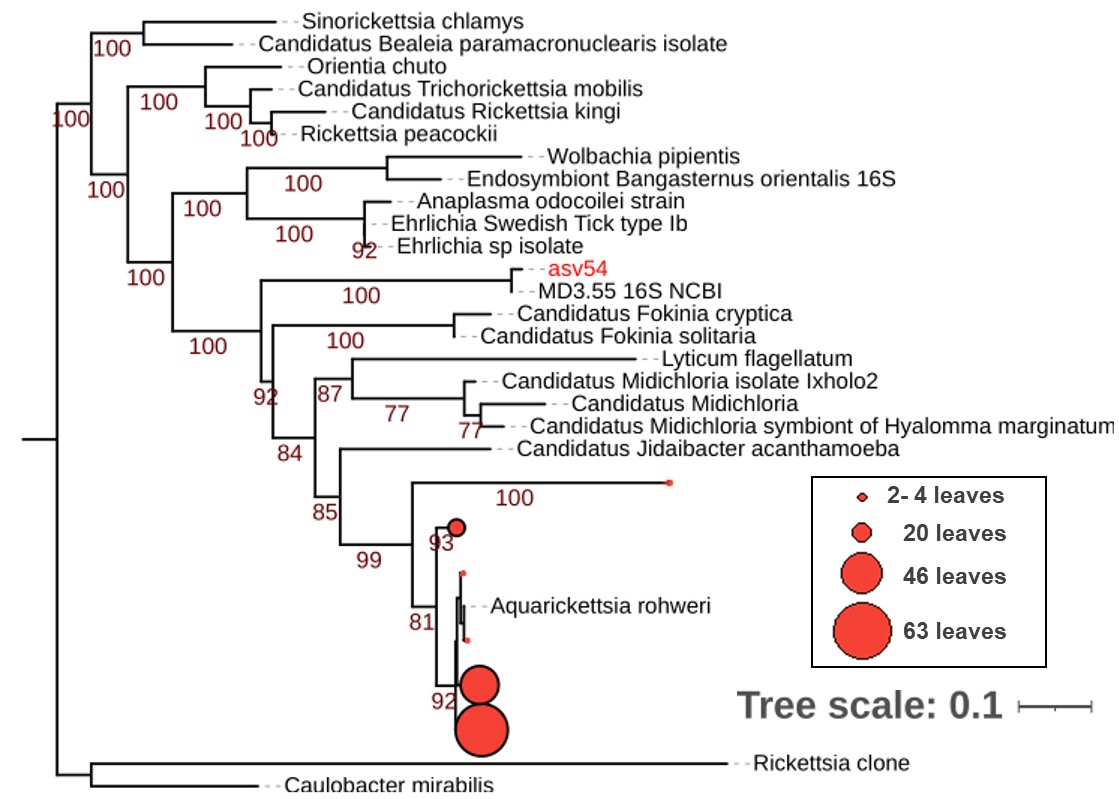

**Figure S5.** Raw mitochondria ASV counts (y-axis) on non-rarefied and non-filtered data in mucus samples collected from the nursery (0) and transplant sites (12). The x-axis shows the tabulated counts for genotypes and background FSW (N: Nursery, BP: Big Pine Shoals, BH: Bahia Honda, LK: Looe Key).
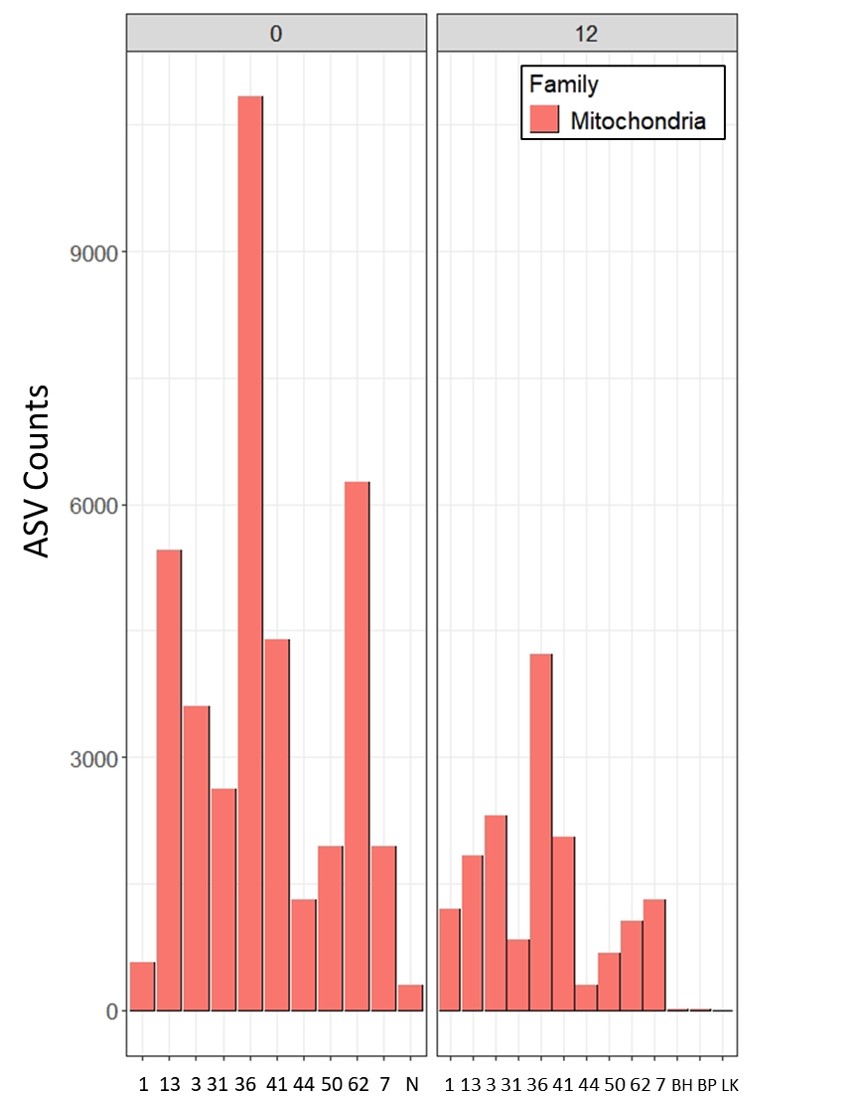

**Table S1**. Coordinates of *Acropora cervicornis* transplant sites and common garden nursery, located off the lower Florida Keys.

| Sites | Latitude | Longitude |
| --- | --- | --- |
| Eastern Dry Rocks | 24.45948 | -81.84412 |
| Marker 32 | 24.47408 | -81.7426 |
| Western Sambo | 24.48268 | -81.70334 |
| E. Sambo | 24.49305 | -81.65961 |
| Maryland Shoals | 24.51036 | -81.56991 |
| Dave's Ledge | 24.53051 | -81.48734 |
| Looe Key | 24.54667 | -81.40214 |
| Nursery | 24.56258 | -81.40008 |
| Big Pine Shoals | 24.56867 | -81.32655 |
| Bahia Honda | 24.58908 | -81.24203 |

**Table S2.** Successfully amplified and filtered (≥ 5,000 reads) sample replicates for each genotype. In the nursery, thirty replicates of each genotype were initially sampled along with three filtered seawater samples (FSW). For T_12_ samples were taken from surviving replicates.

|  | **T0** | | | **T12** | | |
| --- | --- | --- | --- | --- | --- | --- |
| Type (Genotype or FSW) | Amplified/filtered replicates | Initial replicates | Site | Amplified/filtered replicates | Amplified samples per site | Total surviving replicates |
| G3 | 18 | 30 | Nursery | 11 | LK(1), ES(3), DL(1), MR32(2), EDR(2), BH(1), BP(1) | 23 |
| G7 | 7 | 30 | Nursery | 7 | LK(1), ES(2), WS(1), MS(1), EDR(1), BH(1) | 22 |
| G36 | 12 | 30 | Nursery | 7 | ES(3), MR32(1), EDR(2), LK(1) | 26 |
| G41 | 7 | 30 | Nursery | 7 | ES(2), WS(1), MR32(2), LK(1), EDR(1) | 16 |
| G1 | 4 | 30 | Nursery | 10 | BP(2), BH(2), ES(2), MS(1), MR32(1), EDR(1), LK(1) | 18 |
| G13 | 13 | 30 | Nursery | 9 | LK(2),ES(3),DL(1), MS(2), MR32(1) | 21 |
| G31 | 9 | 30 | Nursery | 4 | DL(1), MR32(1), LK(1), BH(1) | 22 |
| G44 | 15 | 30 | Nursery | 9 | BP(2), BH(1), ES(2), MS(1), MR32(1), EDR(1), LK(1) | 22 |
| G50 | 8 | 30 | Nursery | 8 | ES(2), LK(1), MS(2), MR32(1), EDR(1), BH(1) | 23 |
| G62 | 10 | 30 | Nursery | 7 | ES(2), MS(1), MR32(1), EDR(1), BH(1), BP(1) | 16 |
| FSW | 3 | 3 | Nursery |  |  |  |
| FSW |  |  |  | 2 | BH |  |
| FSW |  |  |  | 2 | BP |  |
| FSW |  |  |  | 1 | LK |  |

**Table S3**. Summary of read filtering in the DADA2 ASV pipeline in all samples prior to
data analysis.
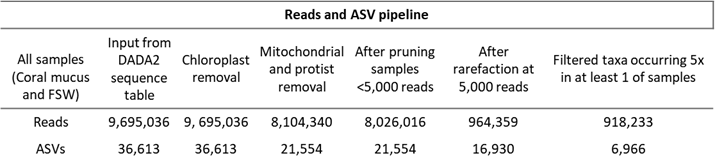

**Table S4.** Alpha-diversity statistical testing output. A Wilcoxon rank sum test was used to conduct a pairwise comparison of epibiome alpha diversity at T_0_ and T_12_ in *A. cervicornis* only. Kruskal-Wallis Rank sum was used to assess differences in epibiome alpha-diversity between the 9 sites.

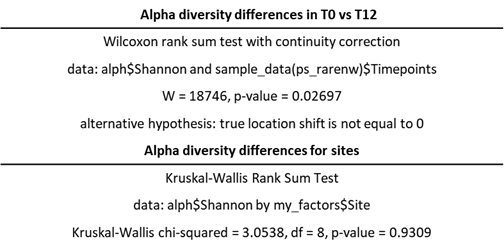

**Table S5**. Beta-diversity statistical output using non-parametric tests on pairwise comparisons. Unevenly dispersed groups were tested using ANOSIM. Groups with even dispersion were tested using PERMANOVA.

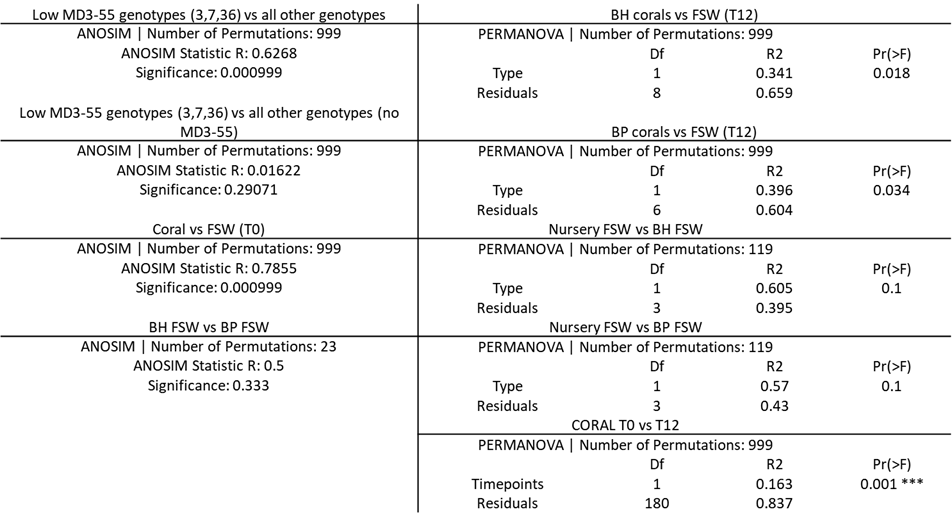

**Table S6.** ANOVA and Tukey HSD output. Differences in alpha-diversity between genotypes were assessed by analysis of variance (ANOVA), followed by a Tukey HSD post-hoc test (95% family-wise confidence level). (* p ≤ 0.05, ** p ≤ 0.01, *** p ≤ 0.001)

A. ***cervicornis* genotypes only**

model aov(formula = alph$Shannon ~ my_factors$Genotype)

Tukey multiple comparisons of means

95% family-wise confidence level

Fit: aov(formula = alph$Shannon ~ my_factors$Genotype)

Genotype	diff	lwr	upr	p adj

13-1	0.221458597 -0.449300941 0.89221814 0.9878967

3-1	0.503978891 -0.134522982 1.14248076 0.2600700

31-1	0.072032045 -0.683646606 0.82771070 0.9999996

36-1	0.672944573 -0.018101680 1.36399083 0.0633594

41-1	0.687685734 -0.053866837 1.42923830 0.0944068

44-1	0.002944413 -0.656856823 0.66274565 1.0000000

50-1	0.415333424 -0.302671765 1.13333861 0.6994436

62-1	0.364696370 -0.343385269 1.07277801 0.8207343

7-1	0.671218563 -0.070334008 1.41277113 0.1131158

3-13	0.282520294 -0.272189346 0.83722993 0.8302659

31-13	-0.149426552 -0.835770756 0.53691765 0.9994991

36-13	0.451485976 -0.162975631 1.06594758 0.3594240

41-13	0.466227137 -0.204532402 1.13698667 0.4417494

44-13	-0.218514184 -0.797613324 0.36058496 0.9699709

50-13	0.193874827 -0.450756536 0.83850619 0.9938258

62-13	0.143237773 -0.490321814 0.77679736 0.9993206

7-13	0.449759966 -0.220999572 1.12051950 0.4955685

31-3	-0.431946846 -1.086801525 0.22290783 0.5199351

36-3	0.168965681 -0.410110610 0.74804197 0.9950663

41-3	0.183706842 -0.454795031 0.82220872 0.9955579

44-3	-0.501034479 -1.042441994 0.04037304 0.0959266

50-3	-0.088645467 -0.699641276 0.52235034 0.9999833

62-3	-0.139282521 -0.738585479 0.46002044 0.9991524

7-3	0.167239671 -0.471262202 0.80574154 0.9978280

36-31	0.600912528 -0.105270828 1.30709588 0.1710302

41-31	0.615653689 -0.140024962 1.37133234 0.2202992

44-31	-0.069087632 -0.744726351 0.60655109 0.9999992

50-31	0.343301379 -0.389284083 1.07588684 0.8896835

62-31	0.292664325 -0.430197775 1.01552643 0.9529609

7-31	0.599186518 -0.156492133 1.35486517 0.2541743

41-36	0.014741161 -0.676305092 0.70578741 1.0000000

**44-36	-0.670000160 -1.272480350 -0.06751997 0.0165832

50-36	-0.257611149 -0.923325923 0.40810363 0.9646450

62-36	-0.308248203 -0.963247681 0.34675128 0.8871208

7-36	-0.001726010 -0.692772263 0.68932024 1.0000000

*44-41	-0.684741321 -1.344542556 -0.02494009 0.0350158

50-41	-0.272352309 -0.990357499 0.44565288 0.9689268

62-41	-0.322989363 -1.031071003 0.38509228 0.9049015

7-41	-0.016467171 -0.758019742 0.72508540 1.0000000

50-44	0.412389011 -0.220832046 1.04561007 0.5386808

62-44	0.361751957 -0.260194233 0.98369815 0.6927107

*7-44	0.668274150 0.008472914 1.32807539 0.0443849

62-50	-0.050637054 -0.734018971 0.63274486 1.0000000

7-50	0.255885139 -0.462120051 0.97389033 0.9794650

7-62	0.306522192 -0.401559447 1.01460383 0.9295260
